# *Caenorhabditis* nematodes colonize ephemeral resource patches in neotropical forests

**DOI:** 10.1101/2022.01.13.476254

**Authors:** Solomon A. Sloat, Luke M. Noble, Annalise B. Paaby, Max Bernstein, Audrey Chang, Taniya Kaur, John Yuen, Sophia C. Tintori, Jacqueline L. Jackson, Arielle Martel, Jose A. Salome Correa, Lewis Stevens, Mark Blaxter, Matthew V. Rockman

## Abstract

Factors shaping the distribution and abundance of species include life-history traits, population structure, and stochastic colonization-extinction dynamics. Field studies of model species groups help reveal the roles of these factors. Species of *Caenorhabditis* nematodes are highly divergent at the sequence level but exhibit highly conserved morphological uniformity, and many of these species live in sympatry on microbe-rich patches of rotten material. Here, we use field experiments and large-scale opportunistic collections to investigate species composition, abundance, and colonization efficiency of *Caenorhabditis* in two of the world’s best studied lowland tropical field sites: Barro Colorado Island in Panamá and La Selva in Sarapiquí, Costa Rica. We observed seven species of *Caenorhabditis*, four of them known only from these collections. While these localities contain species from many parts of the phylogeny, both localities were dominated by globally distributed androdiecious species. We found that *Caenorhabditis* were able to colonize baits accessible only by phoresy, preferring to colonize baits making direct contact with the ground. We estimate founder numbers per colonization event to be low.

## INTRODUCTION

*Caenorhabditis* (Osche, 1952) is diverse. High sequence divergence separates even closely related sister species (Dey *et al*. 2012; Ren *et al*. 2018). Many of these species live in sympatry, yet highly conserved morphology makes it largely impossible to distinguish them without the use of molecular tools or mating tests (Sudhaus & Kiontke 2007). The morphological uniformity among species raises questions about their long-term phenotypic stasis, species coexistence, and the niches they occupy. Previous studies of wild populations of *Caenorhabditis* find that they live on microbe-rich patches of decaying fruit and vegetable matter (Felix & Duveau 2012; Felix et al. 2013; Frézal & Félix 2015; Ferrari *et al*. 2017; Schulenberg & Felix 2017; Crombie *et al*. 2019), the stages on which niche partitioning and interspecific competition play out. Stochastic colonization and extinction rates on these ephemeral resources are key parameters in understanding the local coexistence of species (Dubart *et al*. 2019).

Previous efforts to identify a *Caenorhabditis* substrate niche have classified several species as specialists (Kanzaki *et al*. 2018; Li *et al*. 2014; Dayi *et al*. 2021). The majority, however, have no obvious substrate preference. Studies of *Caenorhabditis* microbiomes in both laboratory (Berg, Zhou, and Shapira 2016) and the field (Dirksen *et al*. 2016; F. Zhang *et al*. 2017) suggest that animals regulate the composition of their gut flora on substrates with differing microbial composition. From these data one could hypothesize that species are specialists, occupying niches defined by what they eat. However, it is unclear which microbes are the primary food source of worms in the wild (Schulenburg & Félix 2017). Beyond food, other factors including predators and pathogens along with non-biological sources of variation like humidity and temperature may play a role in determining where *Caenorhabditis* both colonize and proliferate (Félix & Duveau 2012; Crombie *et al*. 2019). Field experiments have repeatedly concluded that *Caenorhabditis* are much more likely to be found in rotting material than in soil (Frézal & Félix 2015; Schulenburg & Félix 2017). Still missing is substantial evidence that *Caenorhabditis* preferentially colonize specific substrates like fruits or flowers. However, one field study found the degree to which a patch is rotting may influence the incidence of species found on those patches (Ferrari *et al*. 2017).

Equally critical to understanding *Caenorhabditis* adaptation to a metapopulation structure is determining modes of dispersal. Two models described by Slatkin (1977) represent the extremes of a theoretical spectrum. In the *propagule pool* model all colonists are derived from a single patch whereas in the *migrant pool* model colonizers come from the metapopulation at large. In *Caenorhabditis*, these models parallel hypothesized modes of dispersal, either by a phoretic host (Woodruff and Phillips 2018; Sudhaus *et al*. 2011; Kiontke 1997; Yoshiga *et al*. 2013) or by forming a semi-mobile ‘seed bank’ crawling towards or waiting (as specialized dauer larvae) for a fresh patch (Cutter 2015). *Caenorhabditis* are routinely collected from invertebrate vectors, but the contrasting modes have yet to be studied quantitatively. These contrasting modes of dispersal may have profound effects on the level of inbreeding and genetic diversity (Li *et al*. 2014). In addition, propagule size may contribute to the evolution of a female-biased sex ratio and the evolution of self-fertile hermaphroditism as a means of generating a population growth advantage and reproductive assurance, respectively (Theologidis *et al*. 2014; Cutter *et al*. 2019; Hamilton 1967; Lo et al. 2021).

To better understand *Caenorhabditis* diversity and the factors that influence it, we performed field surveys and experiments in two of the most intensively studied lowland tropical forests on Earth: Barro Colorado Island (BCI), Panamá, and La Selva, Costa Rica. Barro Colorado Island lies in the center of the Panama Canal on the man-made Lake Gatún. Shortly after its formation, the island was designated a protected nature reserve and has been hosting field research for the last 100 years (Leigh 1999). Likewise, La Selva Biological Field Station in Sarapiquí, Costa Rica, has been a protected research forest for nearly 70 years (McDade *et al*. 1994). We focused our collection efforts on these two localities as they are relatively undisturbed by human activity and their histories of intensive research provide a rich source of information about the local ecology. One nematode metagenetic study previously found *Caenorhabditis* DNA in a soil and leaf litter sample at BCI, but the species were not identified (Porazinka *et al*. 2010). In contrast to the majority of previous work on *Caenorhabditis* in the tropics, which involved transporting substrates out of country and isolating animals from nematode growth medium plates days or weeks later, we isolated and cultured all animals immediately, in the field, potentially reducing sampling biases that favor species that survive transport and grow well on Nematode Growth Medium. One other study used a combination of these approaches (Félix *et al*. 2013).

In total, we collected seven species of *Caenorhabditis*, four of them known only from these collections (including *C. becei* and *C. panamensis*, which we have described previously [Stevens *et al*. 2019]). Each locality was dominated by globally distributed self-fertile species. We assayed several ecological features related to patch accessibility, patch specificity, and cooccurrence of species. Using baits that vary in their accessibility we demonstrate that *Caenorhabditis* are able to colonize baits that are only accessible by phoresy. Further, colonization rate varied significantly with accessibility where baits making direct contact with the ground were preferentially colonized. We found that individual species tended to occur in habitat patches close to other patches of conspecifics, and we use the frequency of uncolonized patches to estimate the number of colonization events per patch. Taken together our data support current models of most *Caenorhabditis* species as habitat generalists whose population biology is strongly influenced by metapopulation dynamics.

## MATERIALS AND METHODS

### Collections

We collected nematodes on BCI in May 2012 (wet season), March 2015 (dry season), and August 2018 (wet season). Schemes for sampling varied within and among sampling sessions as described in the results. In all cases, worms were isolated from substrates and transferred to Nematode Growth Medium (NGM) plates at the BCI field station and identified as *Caenorhabditis* by morphology under a stereomicroscope. In most cases, material from the forest (*e.g*., rotting fruits and flowers) was placed directly onto NGM plates and *Caenorhabditis* worms picked to new plates to establish cultures (Barrière & Félix 2005). These individual patches of organic material are defined as samples in our dataset and were evaluated for the presence of nematodes. For the majority of samples collected in 2018, worms were isolated by Baermann Funnel technique (Baermann 1917). Cultures of *Caenorhabditis* nematodes were transported to New York for species determination. Species were identified by a combination of 18S and ITS2 rDNA sequencing to derive a prediction and then experimental crosses with isolates of known species identity to establish a biological species assignment (Félix *et al*. 2014; Ferrari *et al*. 2017; Stevens *et al*. 2019).

We collected nematodes at La Selva, Costa Rica, in July 2019, by Baermann Funnel. We used two methods to identify *Caenorhabditis* to species. Individual *Caenorhabditis* worms were chopped with razor blades, transferred to Whatman paper (Marek *et al*. 2014), and transported to New York. There the worms were identified to species by ITS2 sequencing. Separately, we established isofemale cultures on NGM plates. These plates were stored at La Selva for six months prior to their transport to New York, where surviving cultures were revived and species identified by test crosses.

Complete collection data are reported in Supplementary File 1.

### Sequencing and assembling the *Caenorhabditis* sp. 57 transcriptome

We generated the *C*. sp. 57 inbred line QG3077 by 28 generations of full-sib mating from isofemale line QG3050. We generated RNA-seq mRNA transcriptome data using a pool of five mixed-stage populations of QG3077, with each population being subjected to a different condition. All worms were grown at 25 °C on 10 cm NGMA plates (for 1 L: 3 g NaCl, 5 g bacto-peptone, 10 g agar, 7 g agarose, 1 mL cholesterol 5 mg/mL in ethanol, 1 mL CaCl_2_ 1 M, 1 mL MgSO_4_ 1 M, 25 mL KPO_4_ 1 M). One population was fed with CemBio strains (Dirksen *et al*. 2020), and the other four were fed with *E. coli* OP50. The conditions for OP50 populations consisted of 1) mixed-stage, 2) starved, 3) Heat-stressed, 4) Cold-stressed. Temperature stress consisted of exposing the worms to either 35 °C or 4 °C for 2 hours followed by a 2-hour recovery prior to RNA extraction. Total RNA was isolated using TriZol following the protocol described in Green and Sambrook (2020). The mRNA library was constructed using the Illumina Stranded mRNA Prep Ligation protocol. The library was sequenced using a NextSeq 500 MidOutput 2X150 for 300 cycles. Paired-end sequences were trimmed with Trim Galore (https://github.com/FelixKrueger/TrimGalore). Trimmed sequences were assembled into a transcriptome using Trinity (Grabherr *et al*. 2013) also running default parameters for paired-end reads. We then generated the longest predicted ORFs using TransDecoder (https://github.com/TransDecoder/TransDecoder) for use in phylogenetic analyses.

### *Sequencing and assembling the Caenorhabditis* sp. 24 *genome*

After thawing isofemale strain QG555, the nematodes were bleached and grown on 90 mm NGMA plates. We harvested nematodes just after starvation and washed using M9 several times to remove *E. coli*. For genomic DNA extraction, the nematode pellets were suspended in 600 μL of Cell Lysis Solution (Qiagen) with 5 μL of proteinase K (20 μg/μL) and incubated overnight at 56°C with shaking. The following day, the lysate was incubated for one hour at 37°C with 10 μL of RNAse A (20 μg/μL) and the proteins were precipitated with 200 μL of protein precipitation solution (Qiagen). After centrifugation, we collected the supernatant in a clean tube and precipitated the genomic DNA using 600 μL of isopropanol. The DNA pellets were washed in 70% ethanol and dried for one hour before being resuspended in 50 μL of DNAse free-water. For RNA extraction, we resuspended 100 μL of nematode pellet in 500 μL of Trizol (5 volumes of Trizol per volume of pelleted nematodes). The Trizol suspension was frozen in liquid nitrogen and then transferred to a 37°C water bath to be thawed completely. This freezing/thawing process was repeated four to five times and the suspension was vortexed for 30 sec and let rest for 30 sec (five cycles). A total of 100 μL chloroform was added and the tubes were shaken vigorously by hand for 15 sec and incubated for 2–3 min at room temperature. After centrifugation (15 min at 13,000 rpm and 4°C), the aqueous (upper) phase containing the RNA was transferred to a new tube and precipitated with 250 μL of isopropanol. The pellets were washed in 70% ethanol and dried for 15–20 min before being resuspended with 50–100 μL of RNAse-free water. An aliquot of each DNA and RNA preparation was run on agarose gel to check their quality and quantitated with Qubit (Thermo Scientific). Two short-insert (insert sizes of 300 and 600 bp, respectively) genomic libraries and a single short-insert (150 bp) RNA library were prepared using Illumina Nextera reagents and sequenced (125 bases, paired-end) on an Illumina HiSeq 4000 at Edinburgh Genomics (Edinburgh, UK). All raw data have been deposited in the relevant International Nucleotide Sequence Database Collaboration (INSDC) databases.

We performed quality control of our genomic and transcriptomic read sets using FastQC (v0.11.9; Andrews and Others 2010) and used fastp (0.20.1; Chen et al. 2018; --*length_required 50*) to remove low-quality bases and Illumina adapter sequence. We generated a preliminary genome assembly using SPAdes (v3.14.1; Bankevich et al. 2012; --*only-assembler --isolate -k 21,33,55,77*) and identified the likely taxonomic origin of each contig by searching against the NCBI nucleotide (nt) database using BLASTN (2.10.1+; Camacho et al. 2009; -*task megablast - max_target_seqs 1 -max_hsps 1 -evalue 1e-25*) or by searching against UniProt Reference Proteomes database using Diamond BLAST (2.0.4; Buchfink, Xie, and Huson 2015; --*max-target-seqs 1 --sensitive --evalue 1e-25*). We also mapped the genomic reads to the genome assembly using bwa mem (0.7.17-r1188; Li 2013). We provided the assembly, the BAM file, and the BLAST and Diamond files to blobtools (1.1.1; Laetsch and Blaxter 2017) to generate taxon-annotated, GC-coverage plots, which we used to identify contaminant contigs. Any read pair that mapped to the contaminant contigs was discarded. Using this filtered read set, we generated a final assembly using SPAdes (--*isolate -k 21,33,55,77,99*). We also generated a transcriptome assembly using Trinity (Trinity-v2.8.5; Haas et al. 2013), which we then used to scaffold the genome assembly using SCUBAT2 (available at https://github.com/GDKO/SCUBAT2). We used numerical metrics and BUSCO (v4.1.4; Simão et al. 2015; -*l nematoda_odb10 -m genome*) to assess assembly quality and biological completeness, respectively. Prior to gene prediction, we generated a species-specific repeat library using RepeatModeler (2.0.1; A. Smit and Hubley 2010; -*engine ncbi*), and combined this library with known Rhabditid repeats from RepBase (Jurka et al. 2005). This repeat library was then used to soft-mask the genome using RepeatMasker (open-4.0.9; A. F. A. Smit, Hubley, and Green 1996; -*xsmall*). We predicted genes in the genome by aligning trimmed transcriptomic data to the genome using STAR (2.7.3a; Dobin et al. 2013; --*twopassMode Basic*) and providing the resulting BAM file to BRAKER2 for gene prediction (2.1.5; Brůna et al. 2021; --*softmasking*). We used BUSCO (-*l nematoda_odb10 -m proteins*) to assess gene set completeness.

### Phylogenetic analysis

We identified a set of orthologous proteins by running BUSCO (Seppey *et al*. 2019) using the nematode_odb10 dataset on each nematode genome found in Table S1. Multisequence fasta files for each ortholog were extracted using busco2fasta (https://github.com/lstevens17/busco2fasta) with the setting -p 0.8, meaning each ortholog was required to be in 80% or 28 of the 36 species. Orthologous sequences were then aligned with MAFFT (Katoh and Standley 2013) and ML gene trees estimated using IQ-TREE (Nguyen et al. 2015), both on default settings. Newick trees were concatenated into a single file and a species tree was estimated using ASTRAL-III (C. Zhang et al. 2018), which uses a coalescent framework. We also generated a species tree using a supermatrix of all concatenated orthologs. To generate the supermatrix we used TrimAl (Capella-Gutiérrez *et al*. 2009) to remove poorly aligned regions using the settings -gt 0.8 -st 0.001 -resoverlap 0.75 -seqoverlap 80. Sequences were subsequently concatenated using catfasta2phyml (https://github.com/nylander/catfasta2phyml). A tree was then inferred with IQ-TREE using the LG substitution model (Le and Gascuel 2008), modeling the rate variation among sites using a Discrete Gamma model (Yang 1994) with 4 categories. Support was estimated using 1000 ultrafast bootstrap replicates (Hoang *et al*. 2018). We then estimated ASTRAL-III tree branch lengths in units of replacements per site rather than coalescent units using IQ-TREE with the same parameters as the supermatrix analysis while fixing the tree by the output of the ASTRAL-III analysis using the -te setting. All newick trees were visualized using the ITOL web browser (Letunic and Bork 2019).

## RESULTS

### The *Caenorhabditis* faunas of BCI and La Selva

We recovered *Caenorhabditis* nematodes from 225 samples collected on BCI (Figure 1; Supplementary File 1). Additional samples did not contain *Caenorhabditis* or were damaged during processing and shipping. The *Caenorhabditis* isolates derive from opportunistic sampling of rotten fruits, flowers, mushrooms, and leaf litter in 2012 and 2018, from systematic sampling of *Gustavia superba* flowers in 2012, and from several classes of experimental baits in 2015. By DNA barcode sequencing and laboratory mating tests (Sudhaus *et al*. 2011; Félix *et al*. 2014), we assigned the *Caenorhabditis* isolates to six different species, three of which are currently known only from our collections on BCI. These are *C. becei* Stevens 2019, *C. panamensis* Stevens 2019, and *C*. sp. 57. The number of samples yielding each species is shown in Table 1. In total, the 225 samples yielded 260 species observations, as many samples contained multiple *Caenorhabditis* species.

**Figure 1.**
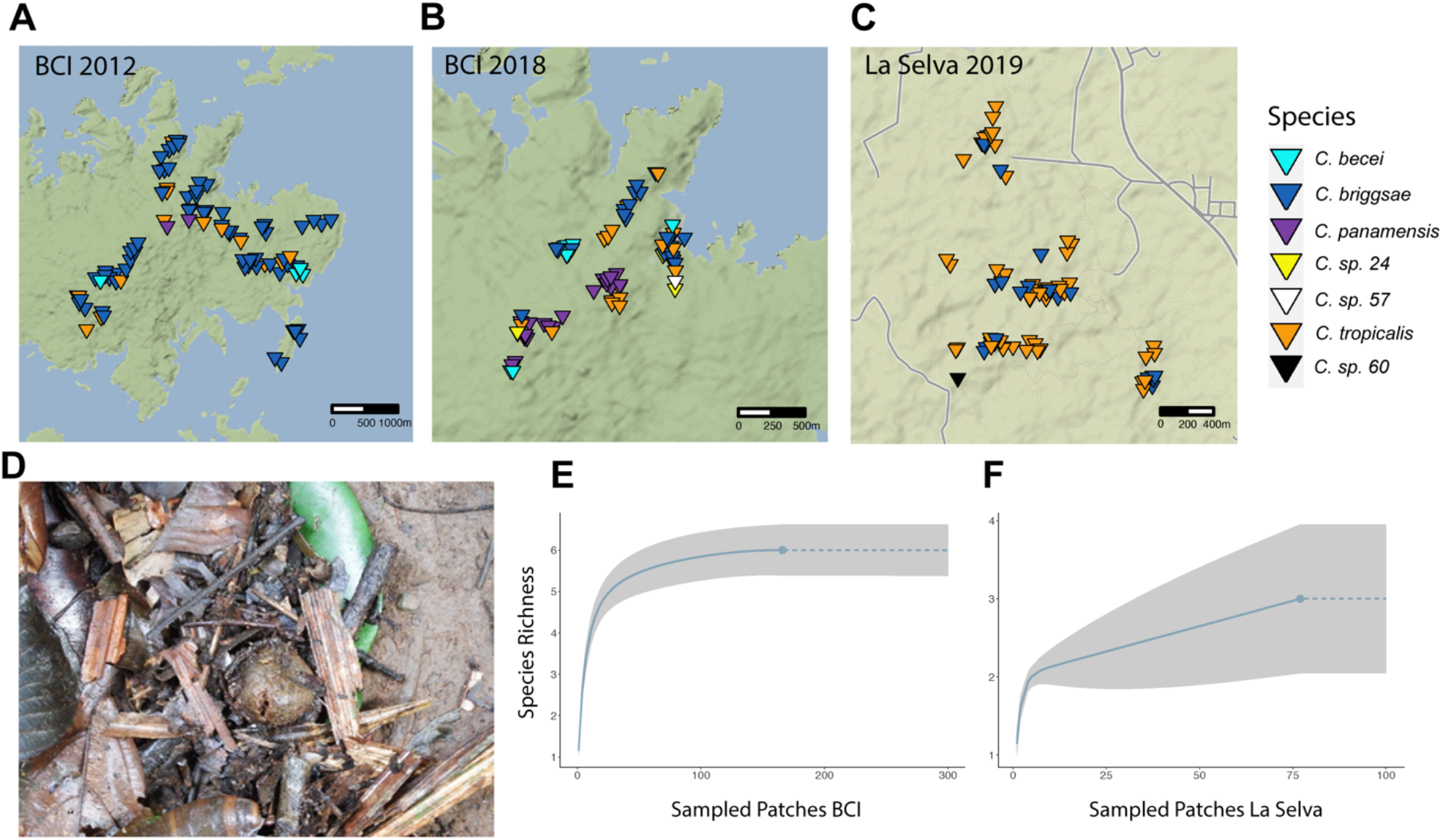
*Caenorhabditis* were collected at two localities: Barro Colorado Island, Panamá and La Selva in Sarapiquí, Costa Rica. **(A-C)** Distribution of species collected from opportunistic sampling from each locality by year. Each marker represents a patch positive for that species. Patches may be plotted multiple times if species co-occurred on the same patch. Patches are jittered to prevent overpotting. **(D)** A fig substrate from which *C*. sp. 57 was isolated. **(E-F)** Rarefaction curve of the chao2 incidence-based estimator for both localities. The solid line represents the predicted species richness the dotted line represents an extrapolation of species richness. The grey area is the 95% confidence interval.

**TABLE 1.**

| Species | Total Positive Samples | 2018 Survey | Range |
| --- | --- | --- | --- |
| *C. briggsae* | 152 | 26 | cosmopolitan |
| *C. tropicalis* | 43 | 15 | pantropical |
| *C. panamensis* | 30 | 11 | endemic |
| *C. becei* | 25 | 10 | endemic |
| *C. sp. 24* | 8 | 6 | neotropical |
| *C. sp. 57* | 2 | 1 | endemic |

To assess the completeness of our survey, we used rarefaction of the chao2 incidence-based estimator (Hsieh, Ma, and Chao 2016; Chao *et al*. 2014), which generated an estimated species richness of 6 ± 0.34 (95% CI) (Figure 1). These data suggest that we have recovered the maximum number of species at BCI, conditional on our sampling strategy. The two most abundant species, *C. briggsae* and *C. tropicalis*, are androdioecious (males and self-fertile hermaphrodites), and their geographic distributions are cosmopolitan and pantropical, respectively. The other species are gonochoristic (males and females). One of these species, *C*. sp. 24, has also been found in French Guiana (Ferrari *et al*. 2017), Mexico, and Southern California (personal observation and Félix 2021).

We successfully recovered *Caenorhabditis* nematodes from 77 samples at La Selva, Costa Rica (Figure 1; Supplementary File 1). These derive from opportunistic sampling of rotten fruits, flowers, mushrooms, and litter in 2019. These samples yielded only 3 different species, one of which is known only from our collections at La Selva (*C*. sp. 60). La Selva differed from BCI in that *C. tropicalis* was most prevalent (present in 55 samples), followed by *C. briggsae* (32 samples). Gonochoristic *C*. sp. 60 was isolated from a single substrate, which contained an estimated thousands of individuals. Rarefaction of the chao2 incidence-based estimator generates a species richness of 3 ± 0.48 (95% CI) (Figure 1). This suggests that the lower number of observed species at La Selva is not due to inadequate sampling given our sampling strategy. We measured substrate temperature for 22 samples that contained *Caenorhabditis;* these ranged from 24.1 to 28.4 °C, with each species averaging 26 °C (Supplementary File 1).

To understand the phylogenetic positions of the undescribed species, we sequenced and assembled a transcriptome for *C*. sp. 57 and a genome of *C*. sp. 24. Using these assemblies and the assemblies of 34 additional *Caenorhabditis* species, we identified 1931 single-copy orthologs that were represented in at least 28 of the 36 species. We estimated the *Caenorhabditis* phylogeny using two approaches. First, we used a coalescent-based approach with individual gene trees as input. Second, we used a maximum likelihood approach using a concatenated alignment of all orthologues as input. The resulting phylogenies (Figure 2) exhibit largely congruent topologies that are consistent with previous analyses (Stevens 2019), differing only in the position of *C. virilis. C*. sp. 24 is closely related to *C. quiockensis* (Stevens 2019) within the Angaria group of spiral-mating species (Sudhaus *et al*. 2011). *C*. sp. 57 is most closely related to *C. monodelphis* (Slos & Sudhaus 2017) and *C. auriculariae* (Tsuda & Futai 1999), which together form the sister group to all other *Caenorhabditis*. From the ITS2 sequence alone *C*. sp. 60 is sister to *C. macrosperma* within the *Japonica* group (NCBI Accession: OL960095). Overall, the species found at BCI and La Selva span the *Caenorhabditis* phylogeny. The two selfers, *C. briggsae* and *C. tropicalis*, are the sole representatives of the *Elegans* group, while three species (*C. becei, C. panamensis, C*. sp. 60) are members of a neotropical-endemic clade within the *Japonica* group.

**Figure 2.**
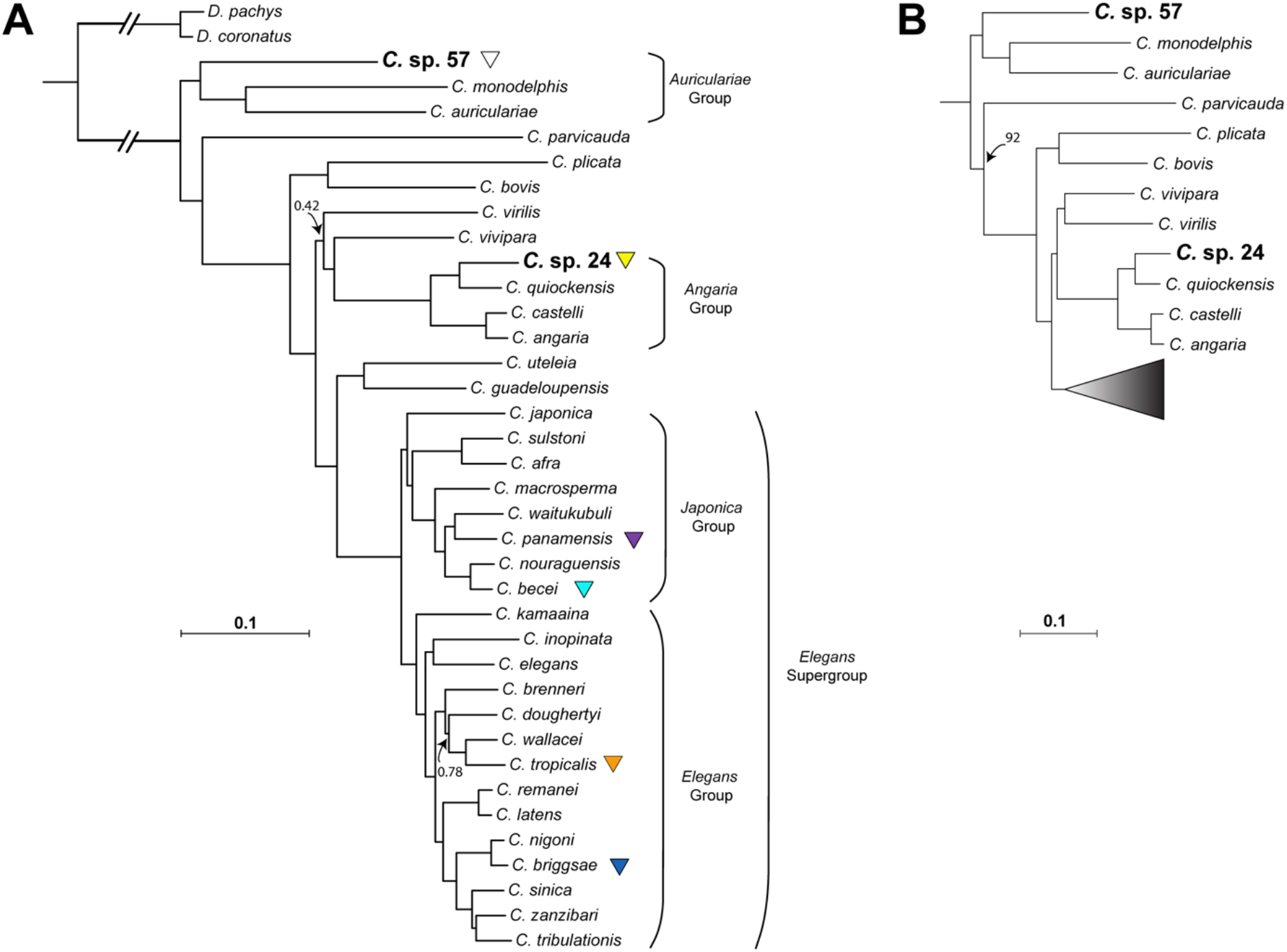
Phylogeny of 36 *Caenorhabditis* species with *D. coronatus* and *D. pachys* forming an outgroup based on 1,931 single-copy orthologs each shared between 80% of the species. **(A)** Phylogeny inferred using a coalescent approach that takes gene trees as input (substitution models for each gene tree selected automatically). Branch lengths in substitutions per site were estimated using the LG substitution model with gamma-distributed rate variation among sites (LG + Γ) while fixing the phylogeny to the coalescent tree topology. Species incorporated into the phylogeny for the first time are bolded. Posterior probabilities are 1.0 unless noted. **(B)** Alternative topology using a supermatrix approach that uses concatenated alignments of all orthologs as input under a LG + Γ model. Bootstrap support is 100 unless noted.

### Substrate specificities

To minimize variation due to differences in sampling technique, we limited our substrate analysis to a dataset of 177 samples collected and processed by a single investigator in August 2018. These samples included a range of rotten fruits, flowers, stems, fungi, and leaf litter. Overall, 94% of the samples yielded nematodes, and 32% (57/177) yielded *Caenorhabditis*. Some samples contained multiple *Caenorhabditis* species, totaling 69 species observations (**Table 1**).

The species abundance ranks match those from the remaining pool of all observations, though the proportions are different, with *C. briggsae* less overwhelmingly dominant. Each of the four most common species was collected from multiple types of fruit and flower. Classifications of the substrates, at high levels (fruit vs other) or lower (*e.g., Spondias mombin* fruit vs. fig), revealed no significant association between *Caenorhabditis* generally or any species specifically and any substrate. Acknowledging the very limited statistical power for most of these tests, we interpret this as evidence that the common species are substrate generalists, colonizing and proliferating in any available habitat patch.

### The spatial patterning of patch occupancy

To understand the spatial patterning of *Caenorhabditis* among habitat patches, we performed hierarchical spatial sampling of a single substrate type, rotten flowers of *Gustavia superba*, in May 2012. We selected four *G. superba* trees spread across the island, and at each we established three well separated 1m^2^ quadrats. Within each quadrat, we sampled four rotten flowers, each at least 10 cm apart. From each flower that yielded *Caenorhabditis*, we established isofemale or isohermaphrodite lines from four or more randomly selected worms from each flower. At one tree only two quadrats were sampled. In total this sampling scheme involved 44 samples of *G. superba* flowers.

Thirty-six of 44 *G. superba* flowers (82%) contained *Caenorhabditis. C. briggsae* was present in every *Caenorhabditis-positive* quadrat at every site, while the other species exhibited strongly patchy distributions over scales of meters (Figure 3). For example, *C. becei* was present in all four flowers in one quadrat at Plot DFT but absent from the flowers in the other two quadrats there. Similarly, *C. tropicalis* was present in three of four flowers in one quadrat at Plot DT but absent from the other two quadrats a few meters away. This patchiness is manifest at larger scales as well: *C. panamensis* was present in all three quadrats at Plot StLT but absent from the other three plots.

**Figure 3.**
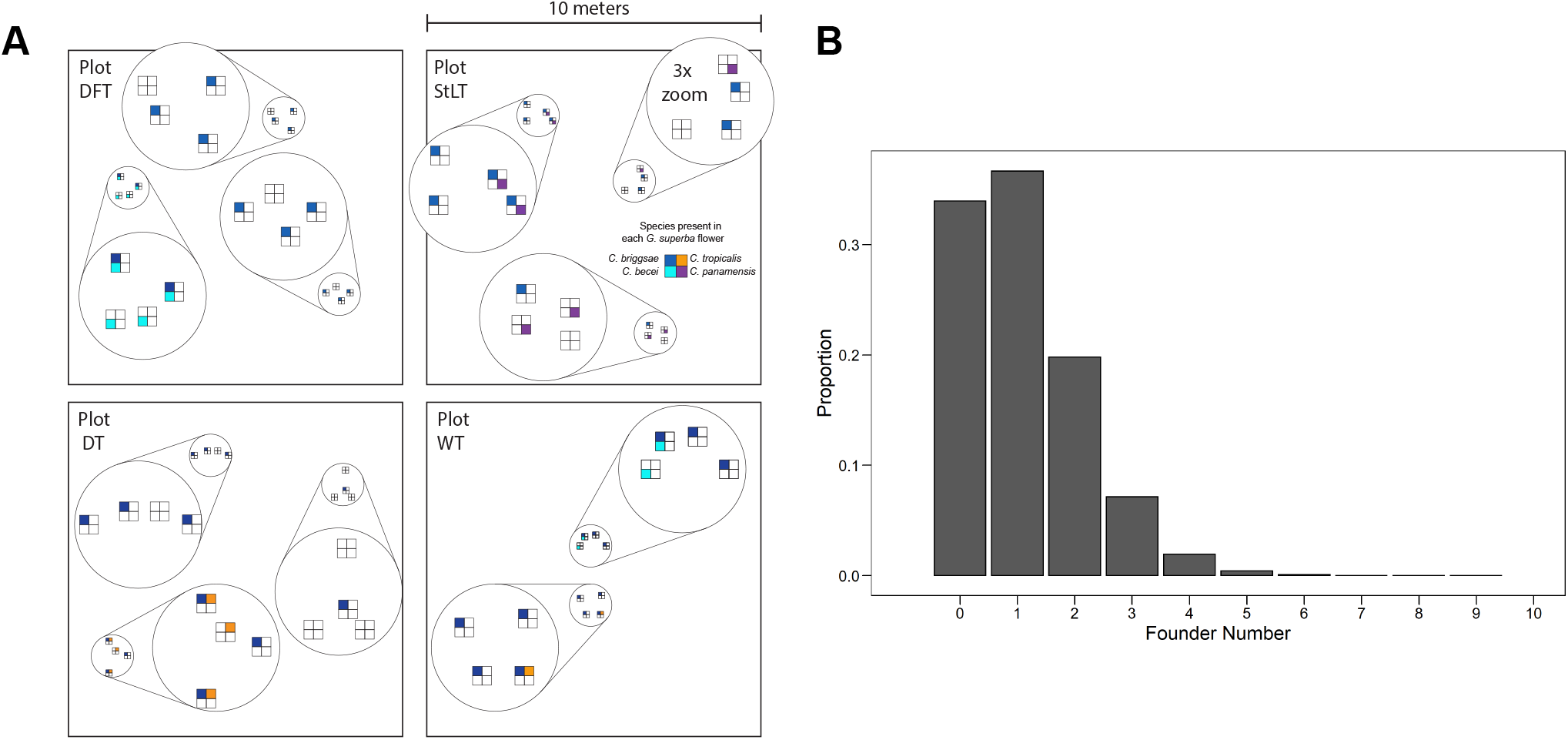
Species are patchily distributed among rotting *Gustavia superba* flowers. **(A)** 10×10 meter plots were systematically sampled at each of four focal trees. At each plot, four flowers were collected from two or three 1-meter quadrats. Each box represents a flower, each color represents the species present on that flower **(B)** The distribution of *C. briggsae* colonization events per flower under a simple Poisson model (mean=1.08).

*C. briggsae* was present in 29 of the 44 *G. superba* flowers (66%). This allows a crude estimate of the number of flowers colonized by *C. briggsae* multiple times. If *C. briggsae* is present ubiquitously and patch colonization is a Poisson process, the absence of *C. briggsae* from 34% of flowers implies a Poisson-distributed number of colonizations per patch with mean 1.08, with 29% of flowers colonized by *C. briggsae* more than once. Thus ~44% of the flowers that contained *C. briggsae* (0.29/0.66) are expected to have had multiple colonizations.

There is no evidence that the presence of one species affects the probability of observing a second species within a sample. For example, *C. briggsae* and *C. tropicalis* are present in 66% and 9% of the 44 samples; the expected co-occurrence under independence is 2.6/44 and we observe co-occurrence of 3/44 samples.

### Colonization patterns among classes of bait

To test how substrate type and accessibility affect rates of colonization by *Caenorhabditis*, we set up arrays consisting of several bait types. At each of seven sites on BCI, we set up a 7-by-7-meter field site with five arrays of baits (four in the corners, one in the center). Each bait array consisted of six agar baits, each bait of a different type (Figure 4), arranged 3×2 with 30 cm spacing between the 6-cm diameter baits. Our experiment as a whole therefore included 210 baits in total. Baits were placed on March 24 2015 and were collected on March 27 2015, at which time a sample of the bait was placed on a seeded plate and the plate was monitored for nematodes twice daily for four days. Twenty-nine of the 210 baits were absent at the time of collection (in cases we observed, eaten by ants and beetles), leaving data for 181 baits for analysis. From each bait that yielded nematodes, we identified *Caenorhabditis* by morphology and established lines. From each *Caenorhabditis-positive* sample we determined the species for at least one line by mating tests.

**Figure 4.**
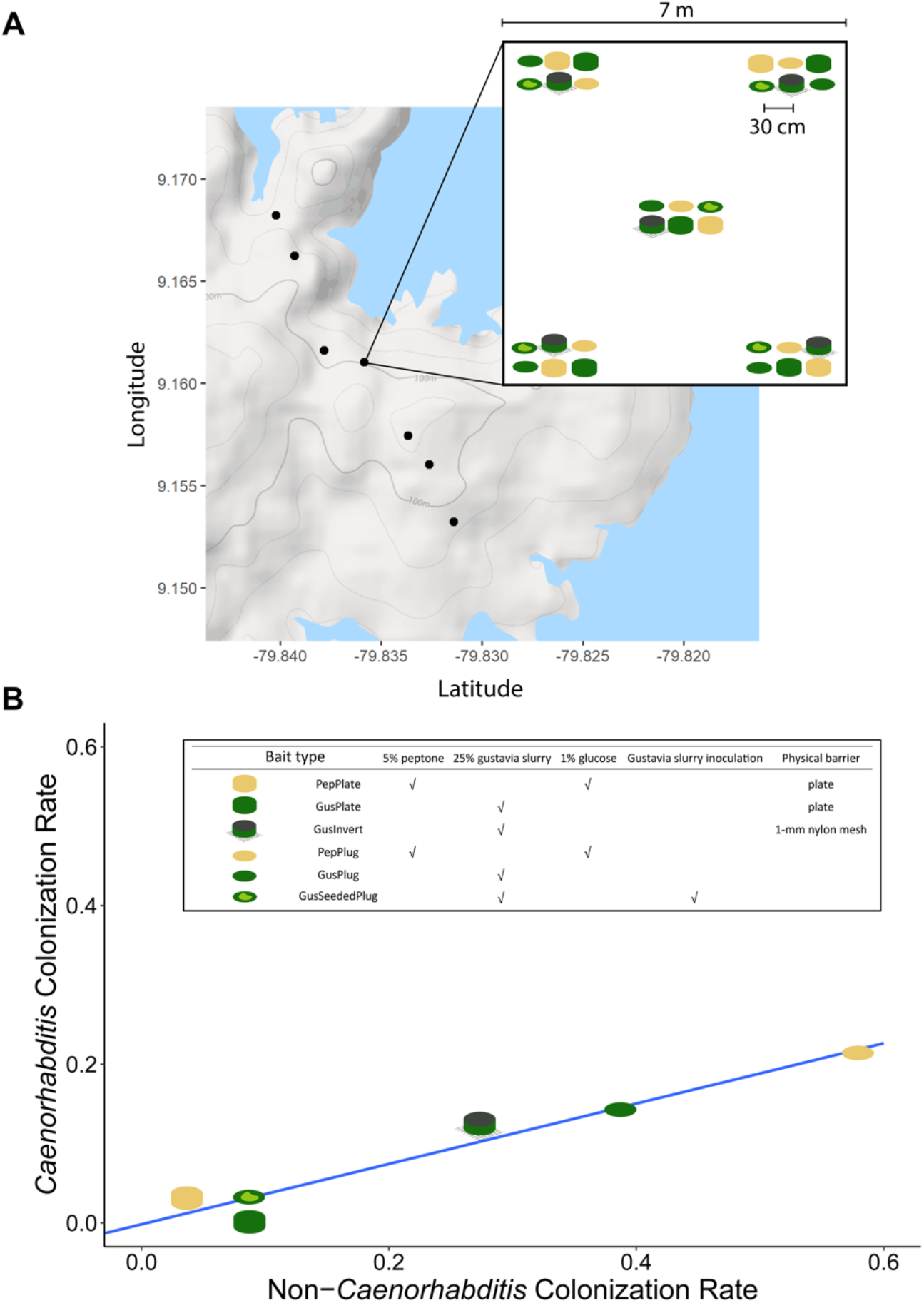
Colonization rates vary in response to bait composition and accessibility. **(A)** Baits were set up at each of seven sites across BCI. Each site consisted of 30 baits arranged in groups of six in the corners and center of each site. **(B)** Six types of agar bait differing in recipe and presence of barriers to access, showed different rates of colonization by nematodes Blue line is linear regression of *Caenorhabditis* vs. *Non-Caenorhabditis* colonization rates across bait types. Table inset contains the composition and accessibility of each bait type.

From 181 baits recovered after three days in the forest, we found 56 (31%) colonized by nematodes, including 17 (9%) colonized by *Caenorhabditis* (15 *C. briggsae* and 3 *C. tropicalis*, including one bait with both species). Colonization rates varied significantly by bait type, for worms overall (p < 10^-12^; analysis of deviance from logistic regression), for *Caenorhabditis* generally (p = 0.001), and for *C. briggsae* specifically (p < 10^-4^). *Caenorhabditis* showed a bait-type distribution that does not differ significantly from the distribution of baits colonized only by *non-Caenorhabditis* nematodes (Fisher’s exact test, p = 0.92), though the power of this test is limited by the small size of the data set. Another way to state this is that the probability of *Caenorhabditis* in a bait type is correlated with the probability of only *non-Caenorhabditis* worms in a bait type (*r*^2^ = 0.98, p < 0.001).

The worms preferentially colonized plug baits, which make direct contact with the ground, over plate baits, which are isolated from the ground by plastic. In both plates and plugs, the worms preferentially colonized those with peptone enrichments over those with heat-defaunated *Gustavia superba* flower slurry. And among *Gustavia* plugs, they preferentially colonized those that were not supplemented with raw *Gustavia* slurry.

### Test of colonization by phoresy

We used size-selective exclosures to determine whether colonization requires phoresy on animals of particular sizes. In 2015, we set up arrays of 24 baits in a 6×4 grid, with 1 meter spacing between samples, at each of six locations spread across BCI. The baits consisted of *Gustavia superba* flower slurry, made by homogenizing flowers and water in a kitchen blender and then heating the mixture to defaunate it. Each array of 24 samples included 4 replicates of 6 different treatments. One treatment consisted of slurry deposited directly onto the forest floor. For the other five treatments, the slurry was placed into a plastic cup and access to the slurry was restricted by the nature of the cup lid. The lids had a circular opening with 3.1 cm diameter, which was either totally open or covered with a nylon mesh to restrict access by animals larger than the mesh size. The mesh openings restricted passage to animals smaller than 4 mm, 1 mm, 0.064 mm, or 0.01 mm.

After 5 days in the field, we collected the slurry samples and transferred a small volume (approximately 1 cm^3^) to NGM plates. If worms emerged, we attempted to establish cultures. Surviving cultures were cryopreserved in New York, and species were identified by sequencing and mating tests. One bait was lost, and of the 143 baits that we recovered, we found nematodes in 30, including three species of *Caenorhabditis* and at least ten additional species (**Figure 5**; **Supplementary File 1**). Because some baits were colonized by multiple species, we count 34 species observations overall.

**Figure 5.**
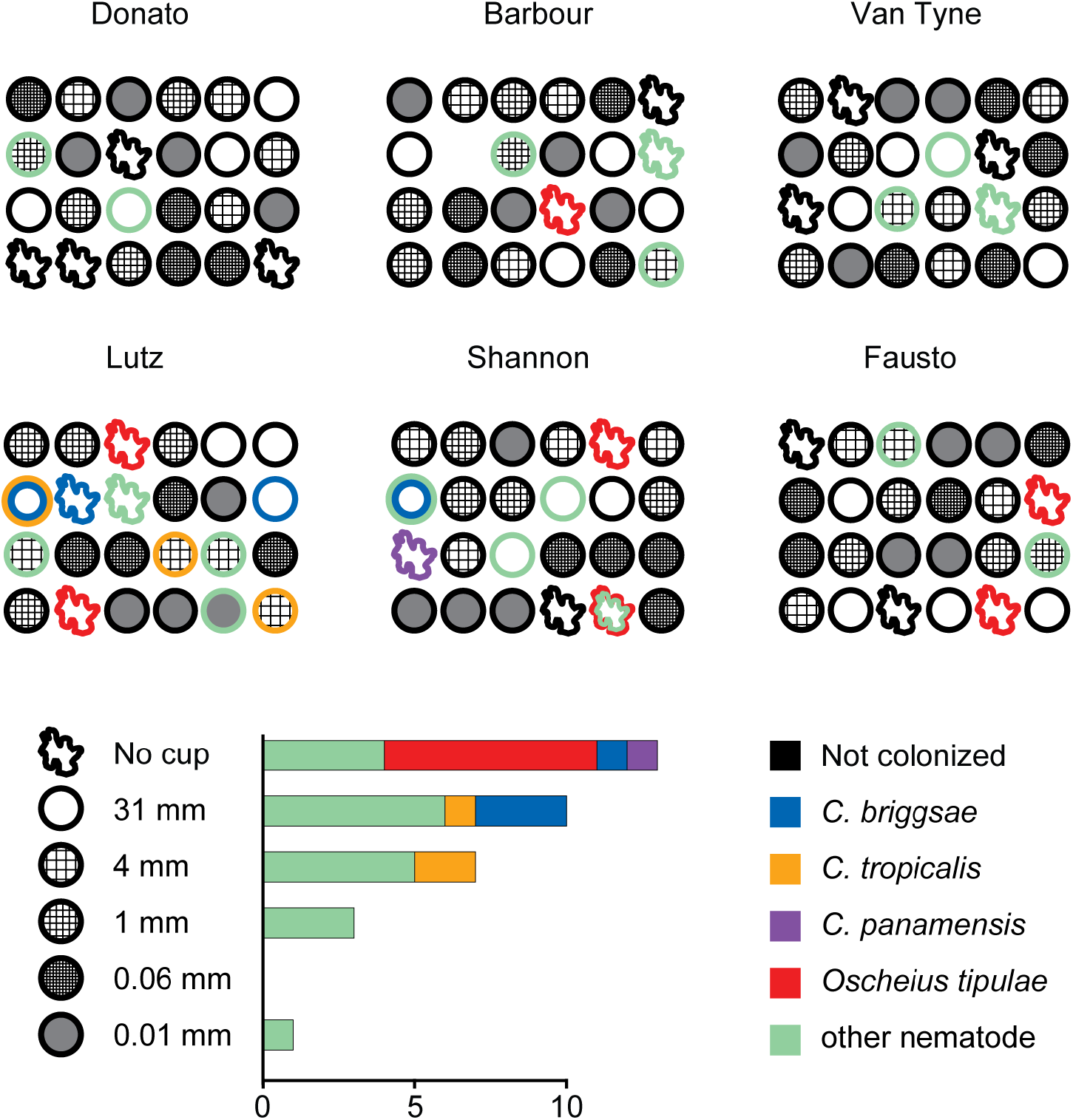
Nematodes colonized 30 baits across six experimental plots, each containing a randomized grid of 4 replicates of each of 6 types of bait differing only in accessibility. Accessibility ranged from no barrier to being accessible via 0.01 mm pores. Colonization varied significantly by bait accessibility. *C. tropicalis* and *C. briggsae* both colonized baits isolated from the environment and accessible only by phoresy while *O. tipulae* was only found to colonize baits making direct contact with the ground.

*C. briggsae* and *C. tropicalis* both colonized baits inside plastic cups, demonstrating that these animals can colonize new substrates by phoresy on other animals. Conversely, *Oscheius tipulae*, which colonized seven baits, only colonized baits that were accessible directly from the soil or leaf litter. We observed substantial heterogeneity among the plots (Figure 4). Bait accessibility significantly affected colonization rates by nematodes generally (p = 5.4×10^-8^; analysis of deviance from logistic regression) and by *Caenorhabditis* specifically (p = 0.007). These analyses treat the bait accessibility as a continuous variable, but analysis with accessibility as unordered levels of a factor yields congruent results. *Caenorhabditis* colonized only the three most accessible classes of bait, suggest that their phoretic hosts were unable or disinclined to pass through mesh with pores of a millimeter or smaller.

## DISCUSSION

Over the past twenty years an increasing community effort to connect the rigorously studied genetic model of *Caenorhabditis* to its natural environment has been fruitful. The catalogue of *Caenorhabditis* species and wild isolates has increased dramatically and along with it the ability to apply population, quantitative, and comparative genomic methods (Stevens *et al*. 2019; Cook *et al*. 2017). Despite these advances, a well-supported model of *Caenorhabditis* population biology is still being formulated. Here, we present a deep sampling of *Caenorhabditis* natural diversity in two of the most extensively studied neotropical field sites, along with a collection of experiments aimed at understanding *Caenorhabditis* in relation to their local metapopulation structure. In total we collected seven species, four of which are only found in these collections (BCI: *C. becei, C. panamensis*, and *C*. sp. 57; La Selva: *C* sp. 60). We estimate that we recovered the total number of species in both field sites accessible to our sampling scheme, which is limited by various factors like time of year, selection of visibly rotting material, nematode isolation method, and proximity of sampling localities to trails. Different sampling schemes would potentially yield different results.

Species from four major clades of *Caenorhabditis* are found in these forests, including representatives of the *Elegans* and *Japonica* groups, the spiral-mating *Angaria* group, and the *Auriculariae* group, which is distantly related to most *Caenorhabditis*. Our findings comport with biogeographic hypotheses about the history of *Caenorhabditis* diversity. In particular, we find three species that are part of a neotropical-endemic clade within the *Japonica* group. Species in this group can be locally abundant in neotropical forests, but their geographic ranges appear to be quite narrow. Each species is known only from a single region, with no overlap among the species in this group found at La Selva, BCI, French Guiana, or Dominica (Stevens *et al*. 2019; Félix 2021). Most parts of the neotropics have not yet been surveyed for *Caenorhabditis*, and we infer that many *Japonica-group* species remain to be discovered there. Conversely, *Elegans* group species are represented exclusively by two widely distributed androdioecious species. Endemic gonochoristic *Elegans-group* species, which are quite numerous in east Asia and Australia, appear to be absent from the neotropics.

Common species at BCI appear to be substrate generalists. Rotten *Gustavia superba* flowers were often occupied by *Caenorhabditis*. We hypothesized that a specific microbial environment on the substrate was preferred by the worms. Our bait preference data suggest that this microbial environment requires conditions that we did not successfully replicate with fresh flower slurry. *Caenorhabditis* preferred baits supplemented with the general microbial growth medium peptone over the *Gustavia* slurry. Moreover, we found that nematode colonization rate was significantly correlated with *Caenorhabditis* colonization rate, suggesting that these worm communities are substrate generalists. This might also suggest that absolute quantity of microbial food is an important factor in determining where *Caenorhabditis* both colonize and proliferate. This conclusion is consistent with our opportunistic sampling data which found no associations between any substrate type and incidence of *Caenorhabditis*. Few field studies have looked at substrate preference specifically. Ferrari *et al*. (2017) found the incidence of *Caenorhabditis* on fresh fruit (citrus) baits to be enriched when compared to non-*Caenorhabditis* nematodes, while Crombie *et al*. (2019) concluded that they observed no substrate specificity between *Caenorhabditis* species and their opportunistically sampled substrates. Future studies would best be served by measuring the response of *Caenorhabditis* incidence to a larger variety of substrate baits and their microbial composition as well as the absolute quantity of microbes on those baits in order to delineate these factors.

Our data supports current models of *Caenorhabditis* modes of dispersal through the use of phoresy in colonization. In our exclosure experiment, *Caenorhabditis* colonized baits that were directly accessible from the ground, isolated from the ground in a cup, and isolated in a cup and further blocked by mesh with openings of 4 mm or greater. Baits isolated by mesh with openings of 1-mm or smaller were not colonized. In contrast, *Oscheius tipulae* only colonized baits making direct contact with the ground (although there is limited evidence for phoresy in *O. tipulae* (De Luca *et al*. 2019)). One hypothesis is that phoretic vectors are colonizing fresh fruit and flower substrates while they are still on the plant, as is the case in some specialist species like *C. inopinata* (Kanzaki *et al*. 2018). Our data confirm that phoretic vectors are mediating the colonization of substrates on the ground, leaving the possibility that animals are also colonizing substrates in trees for future testing.

Species were unevenly distributed over time and geography. There were year-to-year changes in the species collected at various localities around BCI. For example, in 2012 collections at tree DFT yielded *C. briggsae* and *C. becei*, but in 2015 collections at that same tree yielded only *C. tropicalis*. One model is that habitat patches are colonized randomly from the local species pool, as suggested by the patchy species distribution of *G. superba* flower occupancy. An alternative is that species differences among years illustrate ecological succession at larger scales than the level of an individual substrate and its lifespan. Felix & Duveau (2012) more systematically describe a seasonal succession in the abundance of *C. briggsae* and *C. elegans* in a French orchard, paralleling their finding that *C. briggsae* outcompetes *C. elegans* at higher temperatures in the lab. Future analyses of neotropical localities over time will better reveal the spatiotemporal dynamics of the species that coexist there.

Species in our spatial sampling data set appeared to differ in their distributions across sampling sites and quadrats. *C. briggsae* was present in every quadrat at every focal tree sampling site while other species had a patchier distribution over a scale of meters and at scales between focal tree sampling sites. These patterns could indicate differences in colonization efficiency and differences in the scale of dispersal between species which might be picked up by a larger dataset. Under the assumption that animals colonize patches independently and randomly, we estimated that about 44% of patches occupied by *C. briggsae* had multiple colonizations. Richaud *et al*. (2018) modeled *C. elegans* founder number using a Poisson distribution given the proportion of genotypes they observed at a given distance between two patches. They varied how they modeled local haplotype frequencies to account for the unknown proportions in the source population and came to a mean number of 3-10 individuals. Our estimate adds growing support to the hypothesis that founder numbers are low across *Caenorhabditis* and that their population biology is affected by living in an ephemeral metapopulation structure. In general, more detailed investigations into modes of dispersal will reveal a more complete model. These might include characterization of the phoretic vectors employed by *Caenorhabditis* species and genetic analysis of field-collected individuals at fine spatial and temporal scales. Using these data, one could construct a dispersion kernel to understand dynamics and distance of colonization and estimate founder number while minimizing assumptions. Understanding modes of dispersal is crucial to understanding patterns of diversity, inbreeding, and selective pressures that metapopulation structure imposes on traits like selfing and sex ratio.

Our data join with comparable field studies in tropical lowland sites in French Guiana and Hawaii to suggest that androdioecious species not only have larger global ranges than dioecious relatives but are also locally dominant (Table 2). Our collection efforts identify *C. briggsae* as the predominant species at BCI followed by *C. tropicalis*, as in lowland Hawaii (Crombie *et al*. 2019). At La Selva *C. tropicalis* is the most abundant with the sole dioecious isolate being *C*. sp. 60. The largest contrast is Nouragues, French Guiana, where *C. tropicalis* predominates among the androdioecious species but the gonochoristic *C. nouraguensis* is the most abundant overall (Ferrari *et al*. 2017). Taken together, this suggests that the hypothesized benefits of self-fertile hermaphroditism, including reproductive assurance, population growth advantages, and resistance to Medea elements (Cutter *et al*. 2019; Noble *et al*. 2021), are adaptive at multiple spatial scales. While the molecular details of transitions to self-fertile hermaphrodism are well understood (Hill *et al*. 2006; Baldi *et al*. 2009; Woodruff *et al*. 2010; Wei *et al*. 2014), more work needs to be done to understand the factors which drive the evolution of the transition and which maintain dioecious and androdioecious species in sympatry.

**TABLE 2.**
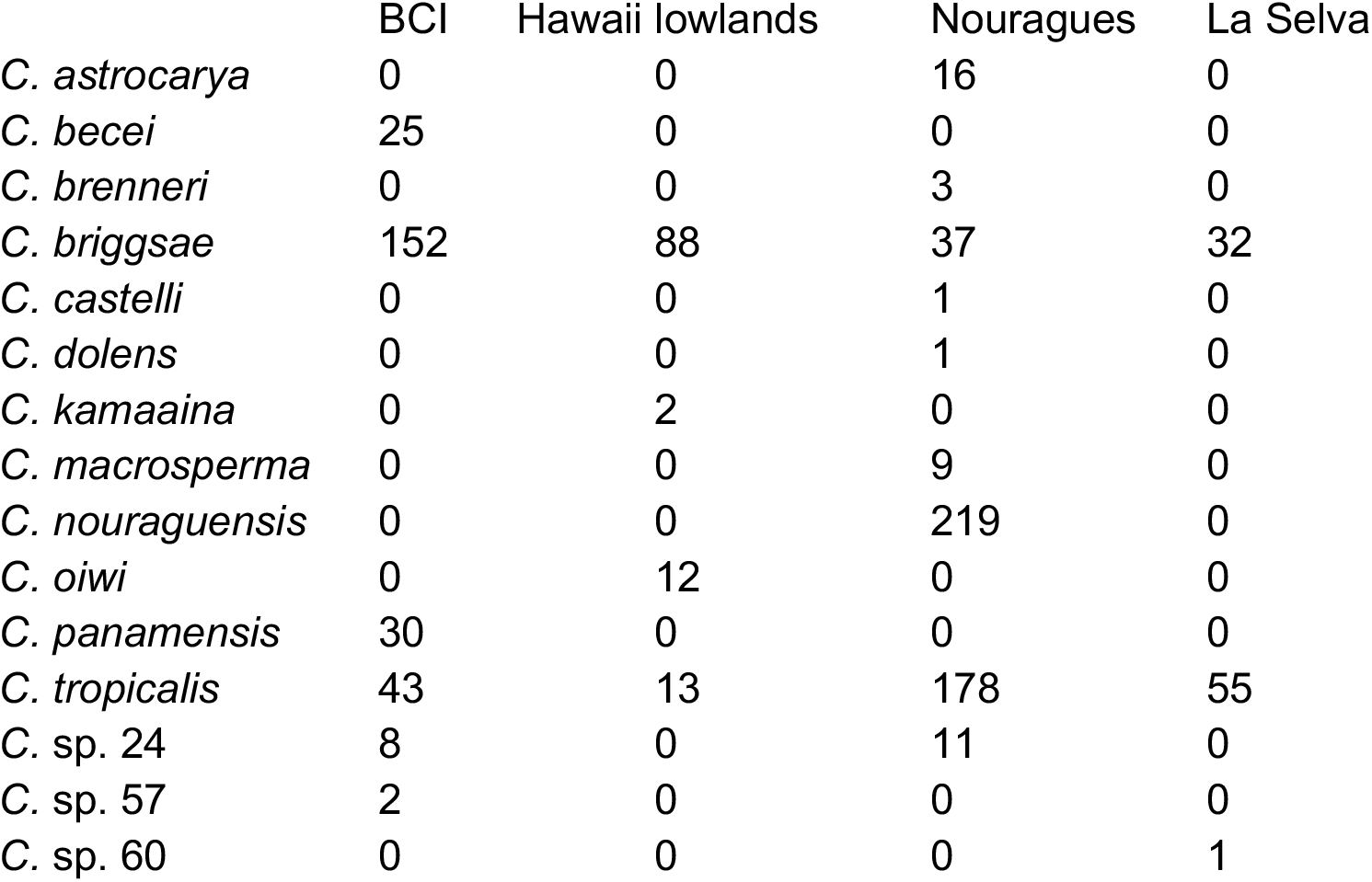

|  | BCI | Hawaii lowlands | Nouragues | La Selva |
| --- | --- | --- | --- | --- |
| *C. astrocarya* | 0 | 0 | 16 | 0 |
| *C. becei* | 25 | 0 | 0 | 0 |
| *C. brenneri* | 0 | 0 | 3 | 0 |
| *C. briggsae* | 152 | 88 | 37 | 32 |
| *C. castelli* | 0 | 0 | 1 | 0 |
| *C. dolens* | 0 | 0 | 1 | 0 |
| *C. kamaaina* | 0 | 2 | 0 | 0 |
| *C. macrosperma* | 0 | 0 | 9 | 0 |
| *C. nouraguensis* | 0 | 0 | 219 | 0 |
| *C. oiwi* | 0 | 12 | 0 | 0 |
| *C. panamensis* | 30 | 0 | 0 | 0 |
| *C. tropicalis* | 43 | 13 | 178 | 55 |
| *C. sp. 24* | 8 | 0 | 11 | 0 |
| *C. sp. 57* | 2 | 0 | 0 | 0 |
| *C. sp. 60* | 0 | 0 | 0 | 1 |

Hawaii Lowlands data are as reported in Crombie *et al*. (2019), including only samples collected in 2017 from elevations below 500m. Nouragues data are as reported in Ferrari *et al*. (2017), representing the count of samples containing each species summed across collections in 2013, 2014, and 2015.

## CONCLUSIONS

Deep sampling of two neotropical sites yielded seven species, four of which are found only in these collections. These collections add growing evidence that self-fertile species not only have larger ranges but are locally dominant in the neotropics. Field experiments support current models of *Caenorhabditis* ecology in which most species are substrate generalists whose population biology is tightly linked to a metapopulation structure. Dauer animals travel from one ephemeral resource to the next on phoretic vectors founding a new population from only a small handful of individuals. These experiments help to inform future work which could more systematically build a model of *Caenorhabditis* metapopulation dynamics which includes species co-occurrence and competition, dispersal dynamics, founding numbers, and the effects of substrate variation and quality.

## Data and code availability

Raw sequencing data and transcriptome assembly for *C*. sp. 24 have been archived under the NCBI study accession PRJEB48807. Raw sequencing data and genome assembly and annotation files for *C*. sp. 57 have been archived under the ENA study accession PRJNA789856.

**Supplementary Table: Phylogenetic Data**

TABLE_S1.xlsx contains the list of all species and accessions to genomic/transcriptomic data used in the phylogenetic analysis.

**Supplementary File: Collection Data**

SupplementaryFile1.xlsx contains all of the collection data reported in the manuscript. Data are provided in a series of sheets corresponding to specific analyses and results, as follows: **BCI.SamplesBySubstrateAllYears**: Each of the 225 rows records one substrate sample that yielded Caenorhabditis nematodes. The species found in each sample are indicated by 1s in the relevant species columns.

**BCI.Opportunistic2012**: Each row represents an isohermaphrodite or isofemale line established from opportunistic collections in 2012.

**BCI.Spatial2012**: Each row represents an isohermaphrodite or isofemale line established from heirarchical spatial sampling of *Gustavia superba* flowers in quadrats around focal trees in 2012.

**BCI.Exclosures2015**: Each row represents a single bait from one of the 24 baits set out at each of six locations in 2015.

**BCI.Exclusures2015Key**: This sheet provides a key to the columns of the BCI.Exclusures2015 sheet, including descriptions of the exclosure types, coordinates of the exclosures, and a summary of the species representation in each type of exclosure.

**BCI.AgarBaits2015**: Each row represents a single bait from one of the 30 set out at each of seven locations in 2015.

**BCI.AgarBaits2015Key**: This sheet provides a key to the columns of the BCI.AgarBaits2015 sheet, including descriptions of the bait types, and coordinates of the field trials.

**BCI.Opportunistic2018**: Each row records one substrate sample that yielded Caenorhabditis nematodes during opportunistic sampling in 2018. The species found in each sample are indicated by 1s in the relevant species columns.

**LaSelva.Opportunistic2019**: Each row records one Caenorhabditis isolate recovered during opportunistic sampling in 2019. Isolates identified by PCR from dried material on Whatman paper have names that start with FTA. Isolates identified from live cultures by mating tests have strain names that start with QG.

**LaSelva.SamplesBySubstrate:** Each row records one substrate sample that yielded Caenorhabditis nematodes during opportunistic sampling in 2019. The species found in each sample are indicated by 1s in the relevant species columns.

## ACKNOWLEDGMENTS

Work in the Barro Colorado Nature Monument was conducted under permits SEX/A-25-12 (2012), SEX/A-28-15 (2015), and SEX/A-55-18 (2018). We gratefully acknowledge the Republic of Panama and the staff of the Smithsonian Tropical Research Institute and Barro Colorado Island Research Station for their assistance. Work at La Selva Research Station was conducted under permits R-033, R-42, and R061-2019-OT-CONAGEBIO. We gratefully acknowledge the Republic of Costa Rica and the staff of the Organization for Tropical Research and La Selva Research Station for their assistance. We thank Christina Zakas and Sarah Rankin for help in the field, Andres Mansisidor, David Riccardi, Patrick Ammerman, Jia Shen, and Aurélien Richaud for help in the lab, and Marie-Anne Félix for helpful comments on the manuscript. This work was supported in part by NIH grants GM089972, GM121828, and GM141906 (MVR) and GM119744 (ABP).

